# Association of brain age with smoking, alcohol consumption, and genetic variants

**DOI:** 10.1101/469924

**Authors:** Kaida Ning, Lu Zhao, Will Matloff, Fengzhu Sun, Arthur W. Toga

## Abstract

The association of the degree of aging based on the whole-brain anatomical characteristics, or brain age, with smoking, alcohol consumption, and individual genetic variants is unclear. Here, we investigated these associations through analyzing data collected for UK Biobank subjects with an age range of 45 to 79 years old. We first trained a statistical model for obtaining relative brain age (RBA), a metric describing a subject’s brain age relative to peers, based on a randomly selected training set subjects (n=2,679). We then applied this model to the evaluation set subjects (n=6,252) and further tested the association of RBA with tobacco smoking, alcohol consumption, and 529,098 genetic variants. We found that daily or almost daily consumption of smoking or alcohol was significantly associated with increased RBA (P<0.05). Interestingly, there was no significant difference of RBA among subjects who smoked occasionally, only tried once or twice, or abstained from smoking. Further, there was no significant difference of RBA among subjects who consumed alcohol 1 to 3 times a month, at special occasions only, or abstained from alcohol consumption. Among the subjects who smoked on most or all days and did not abstain from alcohol drinking, RBA increased by 0.021 years for each addition pack-year of smoking (P<0.05) and by 0.014 years for each additional gram of alcohol consumed (P<0.05). We did not identify individual genetic variation significantly associate with RBA. Further exploration of genetic variation-brain aging association is warranted, where our current genetic association statistics may serve as prior knowledge.

## 1. Introduction

The number of American aged 65 and over is projected to reach 80 million by year 2050 (Ortman et al., 2014). The brain aging process, while associated with structural changes, declined cognitive function, and increased risk of dementia, differs between individuals (Andersen et al., 1999; Jack et al., 2015; Lindenberger, 2014). Therefore, to understand the factors associated with brain aging becomes increasingly important.

It is known that certain lifestyle habits, such as heavy smoking and heavy alcohol drinking are associated with accelerated atrophy in the brain. Compared with non-smokers, smokers have significantly smaller grey matter volume and lower grey matter density in the frontal regions, the occipital lobe, and the temporal lobe. Further, smokers have a significantly greater rate of atrophy in regions that show morphological abnormalities in the early stages of Alzheimer’s disease(Durazzo et al., 2012; Duriez et al., 2014; Gallinat et al., 2006). It has also been reported that patients with alcohol use disorder show decreased regional grey and white matter volumes in the medial-prefrontal and orbitofrontal cortices. The loss of brain gray and white matter volume accelerates with aging in chronic alcoholics (Asensio et al., 2016; Pfefferbaum et al., 1992).

On the other hand, studies have shown that nicotine, a compound contained in tobacco, may improve attention and other cognitive functions in human subjects (Ettinger et al., 2009; Gold et al., 2012). It is also known that drinking wine can be beneficial to the cardiovascular system, which is related to brain health (Almeida et al., 2008; Gianaros et al., 2006; Kappus et al., 2016). Given both the detrimental and potential beneficial effects smoking and alcohol consumption have on the brain, the association of brain aging with smoking and alcohol consumption, especially when the morphology of all the brain regions are considered, remains a subject of investigation.

Besides lifestyle habits, genetic is also associated with brain aging. A recent study analyzed brain imaging data and chronological age (CA) information from twins and suggested that the brain aging process was heritable(Cole et al., 2017b). Currently, the extent to which individual genetic variants are associated with brain aging hasn’t been well studied, except for some conflicting results regarding the association between genetic variation in APOE, a gene associated with Alzheimer’s disease, and brain aging (Cole et al., 2017a; Cole et al., 2018; Lowe et al., 2016).

Recently, researchers have successfully used machine-learning methods to derive a biomarker that is commonly referred to as predicted brain age (PBA) or brain age based on brain imaging data. PBA reflects the degree of aging of the brain based on its anatomical characteristics, as computed based on brain morphology measurements across the entire brain. PBA has been derived and used in several studies, where the mean absolute error between PBA and CA was less than 5 years in adults (Cole and Franke, 2017; Cole et al., 2017b; Franke et al., 2010). Further, it has been shown that advanced brain age is associated with Alzheimer’s disease, objective cognitive impairment, and schizophrenia, etc. (Cole and Franke, 2017; Franke et al., 2013; Franke et al., 2010; Liem et al., 2017; Nenadic et al., 2017).

In this study, we aim to quantify how smoking, alcohol consumption, and genetic variants are associated with brain age. We analyzed the brain imaging data, smoking intensity data, alcohol intake data, as well as genotype data collected for almost 9,000 UK Biobank subjects who were cognitively normal and were of European ancestry. We first trained a model that produces relative brain age (RBA), a metric indicating if a subject’s brain age is older or younger relative to peers, using data for 30% of the subjects. We then applied the trained model to the remaining 70% of the subjects (i.e., the evaluation set) and obtained RBA for those subjects. We found that RBA was associated with various cognitive functions, indicating that RBA captured variations of individual brain aging while adjusting for CA. We further studied the association of RBA with smoking intensity, alcohol consumption, and genetic variants using the evaluation set subjects.

## 2. Materials and methods

### 2.1 Overview of UK Biobank project

The UK Biobank recruited ~500,000 subjects in the United Kingdom(Allen et al., 2014; Sudlow et al., 2015). The participants have provided blood, urine and saliva samples. All participants have been genotyped. The first 10,000 participants scanned as of February 2017 were included in our study (including brain, heart, abdomen, bones and carotid artery). All participants had provided informed consent. The present analyses were conducted under data application number 25641.

### 2.2 Magnetic resonance imaging (MRI) data

Details of the structural brain MRI data, such as imaging hardware and acquisition protocols, are described elsewhere (Miller et al., 2016; Smith et al., 2017). For our analyses, quality controlled structural MRI data was obtained for 9,914 subjects. We excluded 422 (4.3%) subjects with brain and nervous system related illness, including cognitive impairment, neurological disorders or stroke, etc. (see Supplementary Table 1 for the list of diseases based on which subjects were excluded from our analyses). We further excluded 561 subjects with non-European ancestry (according to both self-reported ethnicity and principal component analyses on the genetic data). Brain imaging data of 8,931 subjects were used in our analyses. The age range of these participants is between 45.2 years and 79.4 years.

Brain morphometrics, including volume of cortical, subcortical and white matter regions, thickness and surface area of cortical regions, ventricle size, intracranial volume, etc., were obtained with FreeSurfer 6.0 (Fischl, 2012) based on the T1 MRI brain scans, with the Desikan-Killiany atlas. FreeSurfer is documented and freely available for download online (http://surfer.nmr.mgh.harvard.edu/). Supplementary Table 2 lists the brain morphometric measurements used in our analyses.

### 2.3 Cognitive function

We used the data of cognitive function in its original form, which was collected during the visit for MRI scan. All subjects performed specific tasks as instructed by a computer. To be specific, the Fluid intelligence score indicates the capacity to solve problems that require logic and reasoning ability. It was based on subjects’ performance in identifying the largest number, calculating family relationship, interpolating word, etc. For the prospective memory task, subjects were asked to memorize a command in the middle of the cognitive tests and perform it at the end of the test. In the reaction time test, subjects were asked to press a snap-button when two cards displayed on the computer screen matched. Mean time to correctly identify matches was recorded. In the pairs matching test, subjects were asked to memorize the position of matching pairs of cards. The number of correct pairs identified was recorded. More details of the tasks for assessing cognitive function can be found on the UK Biobank website (http://www.ukbiobank.ac.uk/).

We assessed the statistical significance of the association between RBA and Fluid intelligence score using permutation analyses. To be specific, we first obtained the correlation between Fluid intelligence score and RBA, which was -0.07. We further carried out permutation analyses. In each round of permutation, we permuted the Fluid intelligence scores, and obtained the correlation between RBA and the permutated Fluid intelligence scores (i.e., “permuted correlation”). We did 100,000 permutations and found that none of the “permuted correlations” had an absolute value greater than 0.07. Therefore, we claimed that the correlation between RBA and Fluid intelligence score was significant with a p-value < 0.00001. Other permutation analyses in this manuscript were carried out in a similar way.

### 2.4 Education

We used the information of education qualification collected during the visit for MRI scan. The qualifications are as follows: College or University Degree, A levels or AS levels or equivalent, CSEs or equivalent, NVQ or HND or HNC or equivalent, Other professional qualifications, None of the above. We collapsed the data into two categories as used in the paper by Cox et al. (Cox et al., 2016), indicating whether or not a subject held a college or university degree.

There was no significant association between education and RBA (two-tailed t-test p-value=0.3, Supplementary Figure 11). Therefore, we did not adjust for education when assessing the association of RBA with smoking, alcohol consumption and genetic variants.

### 2.5 Tobacco smoking history and alcohol intake

We used the information of smoking history and alcohol intake status that was collected during the visit for MRI scan. The smoking and alcohol intake frequency categories used in our analyses were as reported in the UK Biobank questionnaire. The smoking pack-years was defined as the number of cigarettes smoked per day/20 multiplied by the number of years of smoking. The alcohol intake amount was calculated as described in the paper by Piumatti et al. (Piumatti et al., 2018). Alcohol consumption per day for a specific type of drink was calculated as the number of drinks consumed per day multiplied by the number of grams of alcohol contained in one drink. The total amount of alcohol consumption per day was the summation of the alcohol amount from all types of drinks. More details can be found on the UK Biobank website (http://www.ukbiobank.ac.uk/).

### 2.6 Genotype data

Details of the genotyping and genotype calling procedures are described elsewhere (UKBiobank, 2015). Quality-controlled genotype data was obtained for 529,098 autosomal SNPs genotyped for 6,195 evaluation set subjects. Our quality control on SNPs ensured that all SNPs had missing rate less than 0.02 and passed Hardy-Weinberg exact test (i.e., Hardy-Weinberg equilibrium p-value >= 1E-6). Quality control on the samples ensured that all subjects had genotyping rate greater than 0.98 and had heterozygosity rate within ±3 standard deviation, had matched reported gender and genetic gender, and were of European ancestry (according to both self-reported ethnicity and genetic ethnicity based on principal component analyses). Related individuals (i.e., kinship coefficient >0.1) were further removed.

### 2.7 Obtaining predicted brain age based on structural MRI data

We first randomly split the brain imaging data of 8,931 subjects into training and evaluation sets. The random sampling ensured that there were no statistically significant differences in the age and gender distributions of the two sets. Further, these two sets had insignificant differences in the smoking and alcohol consumption distributions because of the random sampling (Supplementary Figures 1-2). Our rationale for picking 30% (2,679) of the subjects as the training set and the remaining 70% (6,252) as the evaluation set was to balance the need for accurately training a model to predict brain age and the need for a large number of subjects in the evaluation set for evaluating the association of RBA and the factors of interest.

We then trained a model for predicting brain age based on MRI data using data of the training set subjects. To be specific, we built a linear regression model with Lasso regularization for predicting brain age using R package glmnet (Friedman et al., 2010; R Core Team, 2012). In the model, the chronological age was the response variable, and 403 brain quantitative measures derived using Freesurfer were used as predictors. During model training, the Lasso parameter, lambda, was selected based on an internal cross validation using glmnet. We did not do any pre-selection on the predictors, since the training set sample size was sufficiently large relative to the number of predictors in the model. The mean absolute error (MAE) between PBA and chronological age in the training set was 3.5 years.

After training the model within training set, we applied it to the evaluation set subjects and obtained PBA for those subjects. Since the training and evaluation data did not overlap, the evaluation data also served as an independent validation data for assessing the model’s performance in predicting brain age.

### 2.8 Calculating relative brain age (RBA)

After calculating PBA for each subject, we further calculated a metric that reflected a subject’s PBA relative to peers (i.e., relative brain age or RBA). Due to regression dilution (Hutcheon et al., 2010), the difference between PBA and CA (i.e., PBA-CA) was negatively associated with CA. The older subjects tended to have negative PBACA, while the younger subjects tended to have positive PBA-CA (Supplementary Figure 3). Therefore, when deriving the RBA metric we adjusted for that bias, so that the RBA is directly comparable among subjects at different chronological ages.

To be specific, using the training set data, we built a linear regression model that produced the expected PBA (EPBA) of a subject given the CA while adjusting for gender of that subject (i.e., CA and gender were the predictors and PBA was the response variable).

After training the models for predicting brain age and for further calculating EPBA using the training set data, we applied these models to the evaluation set and calculated the PBA and EPBA for each evaluation set subject. We defined RBA as the difference between PBA and EPBA (i.e., PBA-EPBA). In that way, the mean RBA of all the subjects was zero across all the age ranges. At each age range, there were roughly half of the subjects with positive RBA and half of the subjects with negative RBA (Supplementary Figure 4). A subject with positive RBA has a brain that appears older than those of peers, while a subject with negative RBA has a brain that appears younger.

### 2.9 Quantifying the association of RBA with previous tobacco smoking amount and alcohol intake amount

We quantified the association between previous tobacco smoking amount, alcohol intake amount, and RBA using a two-step regression modeling. We first built a linear regression model using data of 1,316 evaluation set subjects who previous smoked daily or almost daily and did not abstain from drinking alcohol. We then identified subjects with large Cook’s distance as potential influential observations (i.e., subjects with Cook’s distance greater than 3* the mean Cook’s distance of all the subjects). We excluded these influential observations, fitted a second linear regression model, and reported results based on the second regression model. In total, data of 1,231 non-influential observations were used in the second-step regression.

### 2.10 Testing the association between genetic variants and RBA

We used PLINK (Purcell et al., 2007) linear regression model for genotypic test, adjusting for gender and first five genetic principal components of ancestry, to test the association between SNPs and RBA.

We further carried out gene-based and pathway-based association analyses using PASCAL (Lamparter et al., 2016). We included the genes based on the location of the genotyped SNPs (i.e., a SNP located within 2,500-bp region upstream or downstream of a gene is counted as belonging to that gene). The pathways under analyses were collected in the MsigDB database (Subramanian et al., 2005), which is a pathway library combining the results from multiple databases. In total, 18,928 genes and 1,077 pathways from msigDB (Subramanian et al., 2005) were analyzed.

## 3. Results

### 3.1 Computation of predicted brain age (PBA) and relative brain age (RBA)

We trained a model that produced the predicted brain age (PBA) of a subject based on the brain MRI measurements using data of 30% of the UK Biobank subjects. We then applied the trained model to the remaining 70% of the subjects: the evaluation set. The mean absolute error (MAE) between PBA and chronological age (CA) in the evaluation set was 3.9 years. We further obtained relative brain age (RBA) for each subject in the evaluation set (see details in the methods section). Table 1 illustrates the demographic information for the subjects included in the training and evaluation sets. Figure 1 illustrates the relationship between CA and PBA for the subjects included in the evaluation data. We carried out subsequent analyses using data of the evaluation set subjects.

**Table 1.** Demographic information for subjects included in the training and the evaluation data sets.

|  | Number of subjects | Male (%) \| Female (%) | Age (mean [SD], min-max) |
| --- | --- | --- | --- |
| **Training data**<br>(for model training) | 2,679 | 1,274 (48%) \| 1,405 (52%) | 62.6 [7.5], 46.7-79.4 |
| **Evaluation data**<br>(for association analyses) | 6,252 | 2,972 (48%) \| 3,280 (52%) | 62.6 [7.4], 45.2-78.4 |

**Figure 1.**
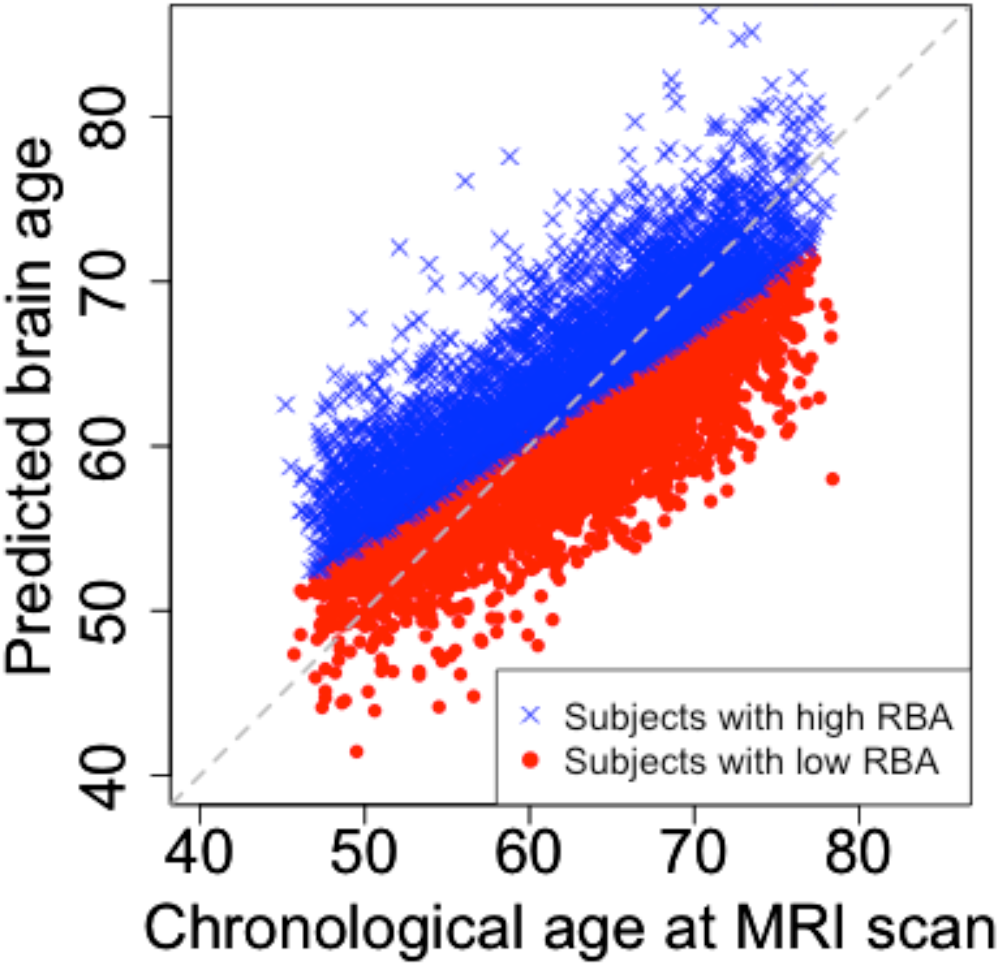
Relationship between chronological age and the predicted brain age. Subjects with higher relative brain age (RBA) are labeled with blue X’s; subjects with lower RBA are labeled with red dots.

### 3.2 Cognitive function is negatively associated with RBA

Subjects who performed better in the cognitive tasks had a lower RBA than that of those who performed worse. For example, Fluid intelligence score was negatively associated with RBA (Spearman’s correlation = -0.07, permutation p-value < 1E-5, see Figure 2). Further, a lower RBA was associated with a better performance in memorizing a specific command and in memorizing the position of matching card pairs, and a lower response time in identifying matching cards. Detailed results on the association between RBA and those cognitive functions are shown in Supplementary Figures 5-9.

**Figure 2.**
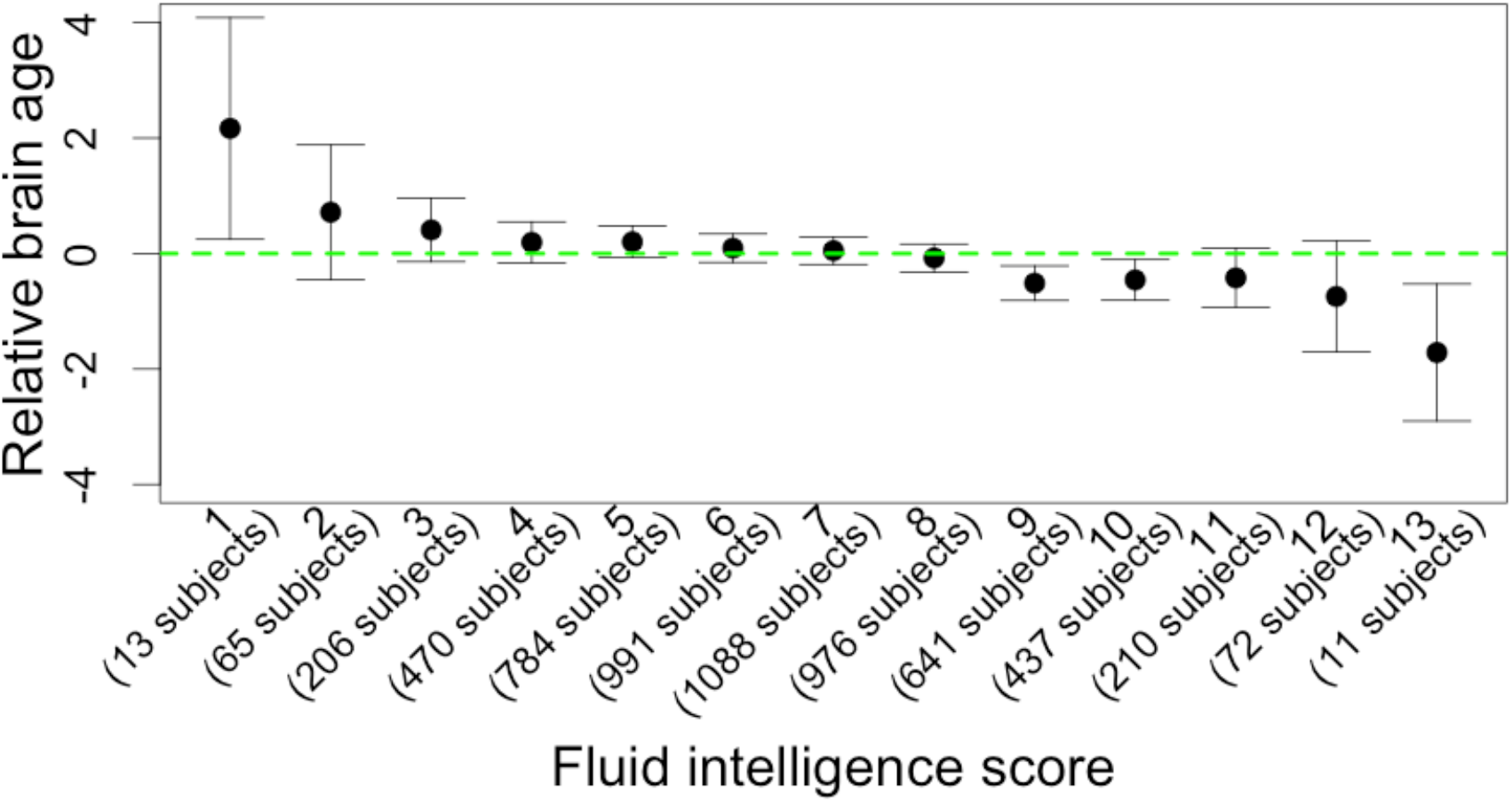
Relationship between Fluid intelligence score and relative brain age (RBA). Subjects with higher Fluid intelligence score have lower RBA.

### 3.3 Previous tobacco smoking and alcohol consumption are significantly associated with RBA

Information of previous tobacco smoking frequency was collected for 6,560 of the evaluation set subjects during the visit for MRI scan. RBA was significantly different among the subjects with different previous tobacco smoking frequency (permutation p-value < 1E-5, Figure 3). Subjects who had smoked on most or all days had the highest average RBA (mean RBA = 0.46 years) compared with those who smoked less frequently. There was no significant difference of RBA among the subjects who smoked occasionally, only tried once or twice, or abstained from smoking.

**Figure 3.**
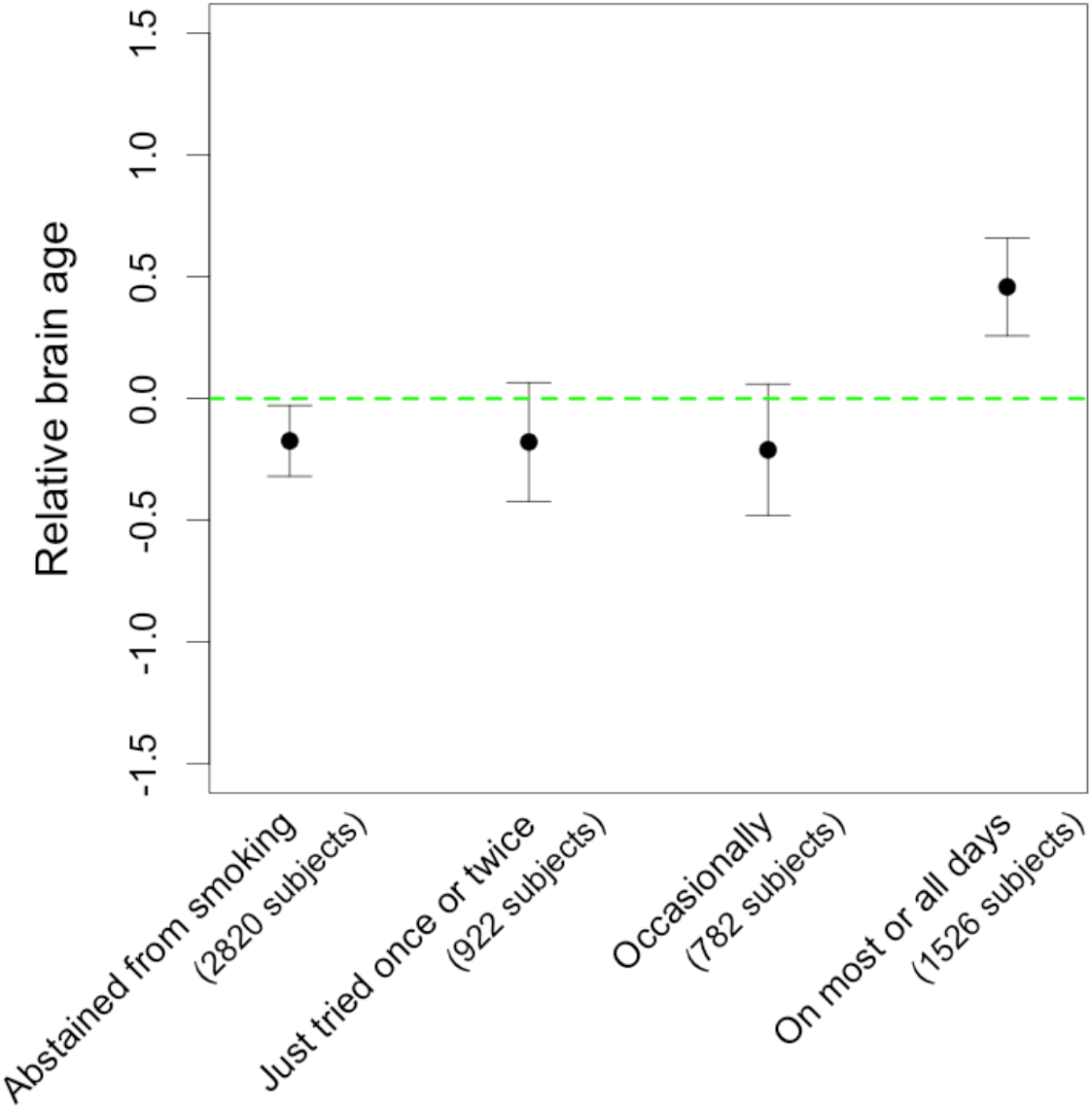
Relationship between previous tobacco smoking frequency and relative brain age.

Information of current alcohol drinking frequency was collected for 6,018 of the evaluation set subjects during the visit for MRI scan. RBA was significantly different among the subjects with different alcohol consumption frequency (permutation p-value = 0.01, Figure 4). Subjects who drank daily or almost daily had the highest average RBA (mean RBA = 0.33 years) compared to those who drank less frequently. The group of subjects who drank at special occasions only had the lowest RBA (mean RBA = -0.33 years), although the RBA difference between those subjects and the subjects who abstained from drinking or the subjects who drank 1~3 times a month was not significant.

**Figure 4.**
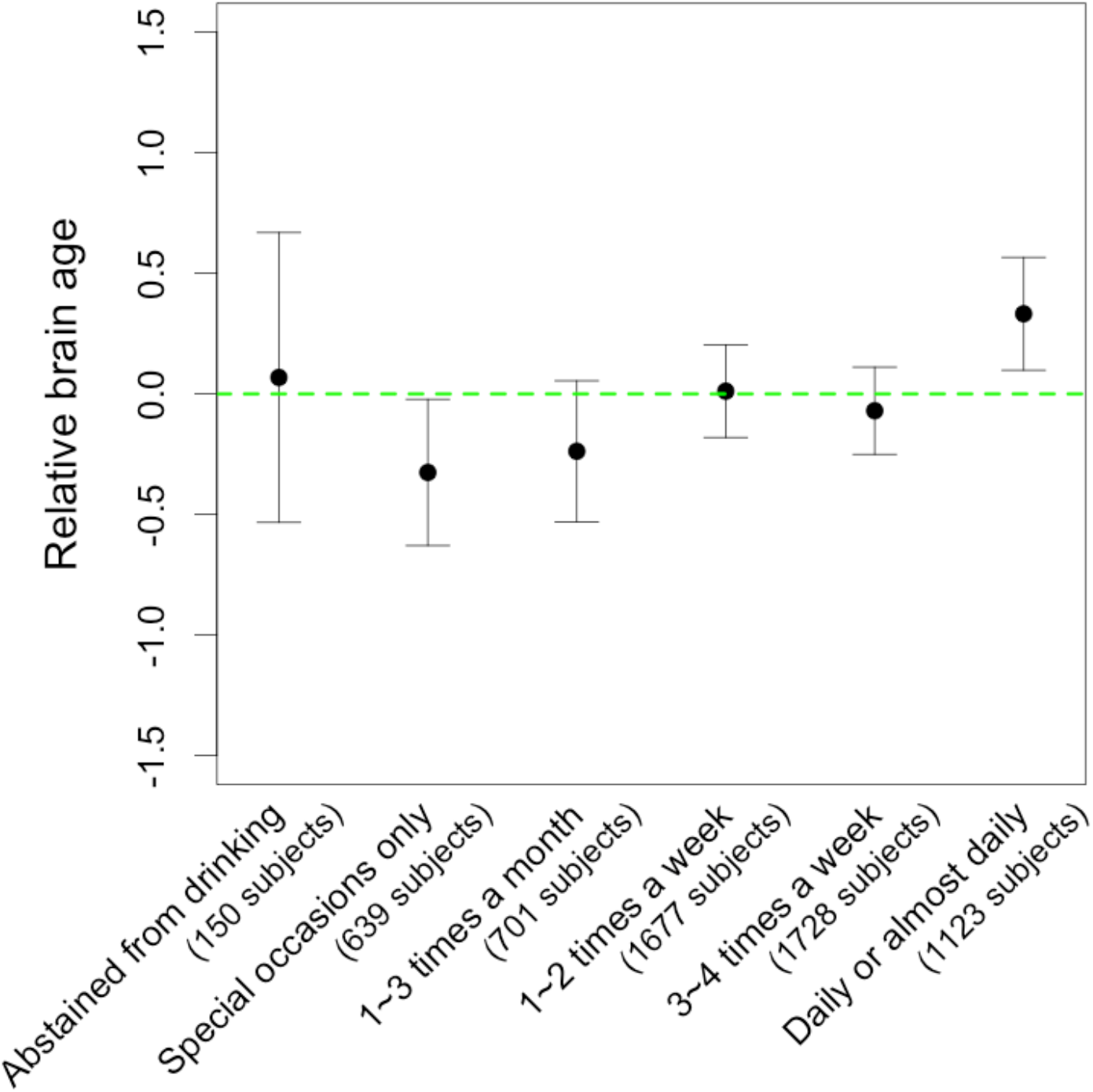
Relationship between alcohol intake frequency and relative brain age.

Smoking and alcohol consumption amount were positively correlated. Among the 1,316 subjects who smoked on most or all days and did not abstain from alcohol, the correlation between the two variables was 0.09 (permutation p-value = 0.001). To quantify the association of RBA with smoking and alcohol consumption, we further built a linear regression model where both smoking and alcohol drinking amount were used as predictors, RBA was the response variable. According to the model, each additional pack-year of smoking was associated with 0.021 years of increased RBA (permutation p-value = 0.013); each additional gram of alcohol consumption per day was associated with 0.014 years of increased RBA (permutation p-value = 0.005). R-squared of the regression model was 0.015. Therefore, high levels of smoking and alcohol consumption were associated with advanced brain age. We also built a regression model with an interaction term between alcohol drinking and smoking. The interaction term was insignificant, indicating that there was insufficient evidence to support the presence of an interaction between alcohol drinking and smoking in affecting RBA.

### 3.4 No significant association identified between single nucleotide polymorphisms and RBA

We looked for single nucleotide polymorphisms (SNP_s_) that were associated with RBA within the evaluation set subjects. The most significant association observed was between SNP rs475675 and RBA (p-value = 2.6E-7). However, the association p-value did not pass the conventional genome wide significance threshold of 5E-8; it was not significant after Bonferroni correction for multiple testing either. The SNP-level RBA association p-values of all the SNPs under analyses are listed in Supplementary Table 3.

We also investigated the association between the dosage of APOE ɛ4 risk allele, a major Alzhiemer’s disease risk factor, and RBA. We found that subjects with two copies of APOE ɛ4 risk alleles had slightly higher RBA than subjects with zero or one copy of risk allele (Supplementary Figure 10). However, the association between APOE risk allele dosage and RBA was insignificant (ANOVA p-value was greater than 0.05).

We further carried out gene-based and pathway-based association analyses to test if certain genetic variants affect brain age in an aggregated way (see details in the Methods section). In total, 18,928 genes and 1,077 pathways from msigDB database (Subramanian et al., 2005) were analyzed. No gene or pathway showed to be significantly associated with RBA after the association p-values were corrected for multiple testing. The gene-level and pathway-level association p-values are listed in Supplementary Table 4 and Supplementary Table 5, respectively.

## 4. Discussion

Here we investigated the association of relative brain age with smoking, alcohol intake and genetic variants through analyzing the data collected for almost 9,000 UK Biobank subjects.

In our analyses, we first calculated PBA of a subject based on structural MRI data and then derived RBA, a metric that describes a subject’s PBA relative to peers. RBA was calculated as the difference between PBA and EPBA (i.e., RBA=PBA-EPBA; see the methods section for details) of a person. As a comparison, in other studies where PBA was derived based on regression model, the difference between PBA and CA (PBA-CA) was used to indicate the brain aging status(Franke et al., 2013; Franke et al., 2010; Nenadic et al., 2017). We used RBA since due to regression dilution, older subjects tend to have negative values of PBA-CA, while younger subjects tend to have positive values of PBA-CA (Supplementary Figure 1). When using RBA, such bias was taken into account. That is, at all age ranges, roughly half of the subjects had positive RBA and half of the subjects had negative RBA.

Our analyses showed that subjects with higher RBA performed worse in various cognitive functions while subjects with lower RBA performed better. A relevant study reported that the biological brain aging accelerated in patients with cognitive impairment than in normal subjects (Liem et al., 2017). Our findings further demonstrated that even among cognitively normal subjects, there was association between advanced brain age and declined cognitive function. We noticed that while the correlation between Fluid intelligence score and RBA was statistically significant, it was not strong. That was due to three main reasons. First, RBA was independent of the chronological age. Therefore, subjects with the same RBA may have a wide range of chronological age, causing large variation of Fluid intelligence score. Second, Fluid intelligence score assesses a subject’s ability to solve new problems, which is one of many cognitive functions the brain carries out. Therefore, the brain function may not be well represented by only Fluid intelligence score. Third, subjects included in our analyses are cognitively healthy. The association between RBA and cognitive function might be relatively weaker within the healthy subjects as compared to a study in which subjects range from cognitively normal, mildly cognitive impaired, and severely cognitively impaired. Nevertheless, the large sample size of our study gave it the statistical power to detect this weak correlation between RBA and Fluid intelligence score.

Our analyses of smoking and RBA indicated that subjects who had smoked on most or all days had a significantly higher RBA compared to subjects who smoked less often. That was consistent with previous studies, which showed significantly greater rate of atrophy in certain regions of the brains of smokers(Durazzo et al., 2012; Duriez et al., 2014; Gallinat et al., 2006). Our data also showed that there was no significant difference of RBA among the subjects who smoked occasionally, only tried once or twice, or abstained from smoking. Previous studies have found that nicotine can help to improve attention and other cognitive functions in human subjects (Ettinger et al., 2009; Gold et al., 2012). It is possible that at a very low amount, the benefit tobacco smoking brings to the brain via nicotine may counteract the detrimental effect it has on the brain. At the same time, we acknowledge that this observation would need to be further validated using an independent data set.

Our analyses of alcohol intake frequency and RBA indicated that subjects who drank daily or almost daily had a significantly higher RBA compared to those who drank less frequently. Our finding was consistent with previous studies, which showed that heavy alcohol consumption was detrimental to the brain(Asensio et al., 2016; Pfefferbaum et al., 1992; Shokri-Kojori et al., 2017). On the other hand, subjects who drank at special occasions only had on average the lowest RBA of all groups of alcohol consumption frequencies. It is known that a small dose of alcohol is associated with a reduced risk of cardiovascular disease, coronary heart disease and stroke(Cleophas, 1999; Corrao et al., 2000; Piumatti et al., 2018; Ronksley et al., 2011). Moreover, cardiovascular health and brain health are related. Researchers have found that cardiovascular risk factors like hypertension and heart disease are associated with increased brain white matter abnormalities and brain atrophy(Almeida et al., 2008; Gianaros et al., 2006; Kappus et al., 2016). Therefore, a small amount of alcohol may be beneficial to brain health through contributing to the cardiovascular health. Our results corroborated the results reported by Gu et al., who showed that light-to-moderate total alcohol intake was associated with larger total brain volume in elderly subjects(Gu et al., 2014).

As for genetic variants, the strongest association between SNP and RBA was 2.6E-7, which was not significant after adjusting for multiple testing. In previous studies, researchers have identified SNPs that showed genome-wide significant association with specific brain morphometrics. For example, SNP rs7294919 (candidate gene TESC) was associated with hippocampal volume; SNP rs945270 (candidate gene KTN1) was associated with putamen volume; SNP rs10784502 (candidate gene HMGA2) was associated with intracranial volume(Hibar et al., 2015; Medland et al., 2014; Stein et al., 2012). It is possible that since brain age was a summary statistic of the morphometrics of multiple brain regions, the associations between SNPs and specific brain regions did not get reflected. Although we did not find any SNP showing genome wide significant association with RBA, the SNP-level RBA association p-values can be used for future meta-analyses, where results from multiple genetic association studies are combined for identifying potentially more significant SNP-phenotype associations.

Several studies had been done to inspect the association between APOE ɛ4 risk allele, a major genetic risk factor for Alzhimer’s disease (AD) (Lambert et al., 2013; Saunders et al., 1993), and brain age. Cole et al. (Cole et al., 2018) looked at the association between APOE ɛ4 status and brain-predicted age difference (PAD) in 669 elderly subjects and reported no association between these two variables. Another study of 30 individuals with Down syndrome reported that APOE genotype did not significantly influence brain-PAD (Cole et al., 2017a). Lowe et al. (Lowe et al., 2016) reported that APOE ε4 status did not have significant association with Brain Age Gap Estimation (BrainAGE) in healthy subjects, patients with AD or mild cognitive impairment. However, they did observe association between BrainAGE changing rates and APOE ε4 carrier status. In our analyses, we found that subjects with two copies of APOE risk alleles had slightly higher RBA than subjects with no risk allele or only one copy of risk allele, although the effect was not statistically significant. Therefore, the effect of APOE risk allele on brain aging is probably not strong within cognitively normal subjects.

Our study has some limitations. First, we used a linear regression model with LASSO to produce PBA based on structural MRI data. More sophisticated statistical models may be built to improve the accuracy of PBA. Also, the combination of structural MRI and other types of brain imaging data (e.g., functional MRI, diffusion-weighted MRI) may help to improve the accuracy of PBA. A more accurate PBA would allow better estimation of RBA. Second, in our study, we investigated the association of brain age with smoking and alcohol intake. Besides smoking and alcohol consumption, various environmental factors may be associated with brain age. For example, physical exercise and meditation had been reported to be associated with lower brain aging level(Luders et al., 2016; Steffener et al., 2016). Therefore, the variation of RBA that can be explained by smoking and alcohol drinking amount was small (as reflected by the small R-squared in the regression model for quantifying the association of RBA with smoking and alcohol drinking amount). More studies can be done to help fully understand the factors associated with brain age. Third, we chose to use pack-years and grams of alcohol intake per day for assessing the smoking and drinking amount. There are alternative measurements for assessing smoking and drinking amount, which may yield slightly different findings(Neuner et al., 2007; Wood et al., 2018). Fourth, it is possible that certain genetic variants that have strong effect on brain age do exist. However, these genetic variants may be missing from the current genotyping platform or they may exist in the form of haplotypes or specific biological pathways and are not detected through current analyses. Fifth, genetic predispositions are known to affect smoking and alcohol drinking behavior. For example, SNPs located in the region of alcohol-metabolizing enzyme genes are significantly associated with alcohol dependence (Hart and Kranzler, 2015). A SNP located in the nicotinic receptor gene is significantly associated with number of cigarettes smoked per day (The_Tobacco_and_Genetics_Consortium, 2010). Therefore, it is possible that genetic variants affect alcohol and nicotine consumption and indirectly affect the RBA. Sixth, a larger sample size would increase the power for identifying SNPs significantly associated with a specific trait. With increased number of UK Biobank subjects for whom both brain imaging and genetic data are available, a future study may reveal SNPs that are significantly associated with brain age.

In sum, we studied the association of brain age with smoking, alcohol consumption, and genetic variants using the data collected for 9,000 cognitively normal UK Biobank subjects. These results provided useful insights into how brain aging is associated with smoking and alcohol consumption. It is still unclear which genetic variants are associated with brain aging. Further studies potentially with even larger sample sizes will be needed to provide a clearer picture of factors associated with brain aging.

## 5. Acknowledgements

This work was supported by grants P41EB015922, U54EB020406 and R01MH094343 of the National Institutes of Health.

We thank Dr. Bo Chen for helpful discussions on the data analyses procedure. We also acknowledge the contributions of members of the UK Biobank coordinating center.

## Supplementary Figures

**Supplementary Figure 1.** Smoking frequency and amount in the evaluation and the training sets.

**Supplementary Figure 2.** Alcohol intake frequency and amount in the evaluation and the training sets.

**Supplementary Figure 3.** Relationship between chronological age and the difference between predicted brain age and chronological age in the evaluation set.

**Supplementary Figure 4.** Relationship between chronological age and relative brain age in the evaluation set.

**Supplementary Figure 5.** Relationship between prospective memory and relative brain age (permutation p-value = 0.01).

**Supplementary Figure 6.** Relationship between the time to correctly identify matches and relative brain age. Red line indicates the regression curve between the two variables (permutation p-value = 0.001).

**Supplementary Figure 7.** Relationship between the number of matches correctly identified and relative brain age (round 1; permutation p-value = 0.005). 202 subjects identified 0 matches, 7 subjects identified 1 match, 1 subject identified 2 matches. Those subjects were grouped together.

**Supplementary Figure 8.** Relationship between the number of matches correctly identified and relative brain age (round 2; permutation p-value = 0.02). 207 subjects identified 0 match, 12 subjects identified 1 match, 1 subject identified 2 matches, 4 subjects identified 3 matches. Those subjects were grouped together.

**Supplementary Figure 9.** Relationship between the number of matches correctly identified and relative brain age (round 3; permutation p-value > 0.05). 3,021 subjects identified 0 match, 7 subjects identified 1 match, 3 subjects identified 3 matches, 1 subject identified 4 matches. Those subjects were grouped together.

**Supplementary Figure 10.** Relationship between education and relative brain age (two tailed t-test p-value > 0.05).

**Supplementary Figure 11.** Relationship between APOE relative brain age (ANOVA ɛ4 risk allele dosage and p-value > 0.05).

## Supplementary Tables

**Supplementary table 1**

List of brain and nervous system related diseases based on which subjects are excluded from the analyses.

**Supplementary table 2**

List of brain measurements used as predictors in the linear regression model.

**Supplementary Table 3**

P-values from the tests for association between each SNP and relative brain age.

**Supplementary Table 4**

P-values from the tests for association between each gene and relative brain age.

**Supplementary Table 5**

P-values from the tests for association between each pathway and relative brain age.

